# *Schizosaccharomyces versatilis* represents a distinct evolutionary lineage

**DOI:** 10.1101/2023.09.04.556188

**Authors:** Graham J Etherington, Elisa Gomez Gil, Wilfried Haerty, Snezhana Oliferenko, Conrad A. Nieduszynski

## Abstract

The fission yeast species *Schizosaccharomyces japonicus* is currently divided into two varieties – *S. japonicus* var. *japonicus* and *S. japonicus* var. *versatilis*. Here we examine the var. *versatilis* isolate CBS5679. The CBS5679 genome shows 88% coding sequence identity to the reference genome of *S. japonicus* var. *japonicus* at the coding sequence level, with phylogenetic analyses suggesting that it has split from the *S. japonicus* lineage 25 million years ago. The CBS5679 genome contains a reciprocal translocation between chromosomes 1 and 2, together with several large inversions. The products of genes linked to the major translocation are associated with “metabolism” and “cellular assembly” ontology terms. We further show that CBS5679 does not generate viable progeny with the reference strain of *S. japonicus*. Although CBS5679 shares closer similarity to the “type” strain of var. *versatilis* as compared to S. *japonicus*, it is not identical to the type strain, suggesting population structure within var. *versatilis*. We recommend that the taxonomic status of *S. japonicus* var. *versatilis* is raised, with it being treated as a separate species, *Schizosaccharomyces versatilis*.

**Take-away:**

- The taxonomic status of *Schizosaccharomyces versatilis* is addressed.
- *S. versatilis* diverged from *S. japonicus* around 25 million years ago.
- *S. versatilis* does not produce viable progeny in crosses with *S. japonicus*.
- *S. versatilis* has a reciprocal translocation between chromosomes 1 and 2.
- The Gene Ontology terms for genes in the translocations are enriched for terms connected to “metabolism” and “cellular assembly”.

## Introduction

Schizosaccharomyces is a genus of symmetrically dividing pill-shaped yeasts within the Taphrinomycotina subdivision of Ascomycota fungi (Kurtzman & Sugiyama, 2015; Liu et al., 2009; Sugiyama, Hosaka, & Suh, 2006). Whilst Saccharomycetes, another class of Ascomycota yeasts, are taxonomically diverse, containing numerous genera (including the model organism *Saccharomyces cerevisiae*), Schizosaccharomycetes contain only one genus – *Schizosaccharomyces*, which is currently subdivided into only six recognised species – *S. pombe, S. octosporus, S. cryophilus, S. osmophilus, S. lindneri* and *S. japonicus* (Brysch Herzberg, Jia, Seidel, Assali, & Du, 2022). *Schizosaccharomyces pombe* is a widely used model organism in diverse fields of molecular cellular biology research, which has provided fundamental insights into the regulation of cell growth and cell cycle, mitosis, meiosis, chromosome biology, RNA processing, epigenetics, cellular organization, polarity establishment and maintenance, cytoskeleton function, etc (Wood et al., 2002; Wood et al., 2012) . More recently, another fission yeast species, *Schizosaccharomyces japonicus*, has emerged both as a part of a powerful composite evolutionary cell biology model system used alongside *S. pombe* (Y. Gu, Yam, & Oliferenko, 2015; Makarova et al., 2016; Makarova et al., 2020; Yam, He, Zhang, Chiam, & Oliferenko, 2011) and as a valuable standalone model organism for the study of processes not apparent or tractable in other yeasts (Aoki et al., 2011; Chapman, Taglini, & Bayne, 2022; Furuya & Niki, 2010, 2012; Gomez-Gil et al., 2019; Y. Gu & Oliferenko, 2019; Kinnaer, Dudin, & Martin, 2019; Lee et al., 2020; Nozaki, Furuya, & Niki, 2018; Papp, Acs-Szabo, Batta, & Miklos, 2021; Pieper, Sprenger, Teis, & Oliferenko, 2020; Rutherford, Harris, Oliferenko, & Wood, 2022; Wang et al., 2021; Yam, Gu, & Oliferenko, 2013).

It has been proposed that *S. japonicus* has two varieties (here denoted as ‘var.’): *S. japonicus* var. *japonicus*, and *S. japonicus* var. *versatilis. S. japonicus* was first isolated in 1911 from Chapel Hill, USA (Brysch Herzberg et al., 2022), although not properly described until 1931 (Yukawa & Maki, 1931). *S. japonicus* var. *versatilis* was isolated in 1945 in Illinois, USA, from fermenting grape must (Wickerham & Duprat, 1945). Another strain of var. *versatilis* was discovered in the slime flux of an elm tree *Ulmus carpinifolia* at the University of California, Davis in 1960 (Phaff, Yoneyama, & CARMO-SOUSA, 1964). That strain has been deposited in the Westerdijk Institute collection as CBS5679. More recently, CBS5679 has been referred to as *S. japonicus* var. *japonicus* in yeast collections (e.g. https://www.dsmz.de/collection/catalogue/details/culture/DSM-70571, https://www.atcc.org/products/24256) and literature (Bouwknegt et al., 2021). In terms of taxonomic assignment, *S. japonicus* var. *versatilis* has a chequered history. Based on a variety of properties (e.g. iodine reaction of mature ascospores, cellular cytochrome spectra, prototrophic selection hybridisation, nDNA/nDNA optical reassociation, and ascospore shape), it has been split from and lumped with var. *japonicus* several times (Johannsen, 1981; Martini, 1991; Sipiczki, Kucsera, Ulaszewski, & Zsolt, 1982; Slooff, 1970; Yarrow, 1984).

Advances in whole genome sequencing and assembly methods have made it possible to sequence and assemble the relatively small genomes of yeast species cheaply and quickly. All known species of fission yeast have now been sequenced (Brysch-Herzberg et al., 2023; Jia et al., 2023; Rhind et al., 2011; Wood et al., 2002). More recently, the availability of low-cost long-read sequencing combined with advances in genome assembly algorithms has allowed for chromosome-scale telomere-to-telomere sequencing to become achievable even for the most complex genomes (Belser et al., 2021; Nurk et al., 2022).

Although genome-wide sequencing data are often generated for genome assembly, other questions can be addressed with the same data types. Recently, whole genome sequencing of the *S. japonicus* strain CBS5679 using long and short-reads was carried out in the course of inquiry into the mechanistic basis of anaerobic growth (Bouwknegt et al., 2021). We noticed that the sequences produced during this experiment differ greatly from those of the reference *S. japonicus* assembly (Rhind et al., 2011; Rutherford et al., 2022).

Here, we assemble the data generated by Bouwknegt (2021) and compare them to the reference assembly of *S. japonicus* and other *Schizosaccharomyces* species.

## Materials and Methods

### Genome assembly and quality control (QC)

We accessed publicly available DNA sequence data for *S. japonicus* CBS5679 generated under BioProject Accession PRJNA698797, which comprised Oxford Nanopore (ONT) long-reads (SRR15295841) and Illumina short-reads (SRR15295840) (Bouwknegt et al., 2021). We used Canu (Koren et al., 2017) to correct and trim the ONT reads and then generated an assembly using reads of >60 kb (resulting in 29,650 reads with an estimated coverage of 42x). We further polished the final assembly with the Illumina data using three rounds of Pilon polishing (Walker et al., 2014). We used BUSCO (v5.3.2) (Manni, Berkeley, Seppey, & Zdobnov, 2021) to identify the number of Ascomycota-specific single-copy orthologs recovered in the assembly. We then carried out repeat masking using a pipeline with RepeatModeler (Flynn et al., 2020) and RepeatMasker (Smit, Hubley, & Green, 2015; Tarailo-Graovac & Chen, 2009) and summarised the results using the ParseRM.pl script from the Parsing-RepeatMasker-Outputs tool (https://github.com/4ureliek/Parsing-RepeatMasker-Outputs).

### Annotation

Liftoff (v1.5.1) (Shumate & Salzberg, 2020) was used to lift over the Ensembl Fungi *S. japonicus* (GCF_000149845.2_SJ5) annotation to CBS5679. Annotation of repeat content was generated using RepeatModeler v2.0.3 (Flynn et al., 2020) and RepeatMasker (v4.1.2-p1) (Smit et al., 2015; Tarailo-Graovac & Chen, 2009). tRNA loci were annotated using tRNAscan-SE (v2.0.12) (Chan, Lin, Mak, & Lowe, 2021). rRNA loci were annotated using barrnap (v0.9) (Seeman, 2018).

### Comparison to S. japonicus

We used Nanopore reads from the type strain (FY16936) of *S. japonicus* (catalogue number ATCC10660) from ENA Project PRJEB63404 and included it in the following analyses. We also included a telomere-to-telomere genome assembly of *S. japonicus* (FY16936; GCA_956476325) to add resolution around the *japonicus*-*versatilis* analyses.

To compare sequence identity at the coding sequence (CDS) level between CBS5679 and *S. japonicus* FY16936, we downloaded the CDS sequences of *S. japonicus* from Ensembl Fungi (database version 109.2) and created a BLAST (v 2.7.1) database from the Nanopore reads of both FY16936 and CBS5679. We then used the Ensembl CDS sequences (4,892 in total) as the query sequences in separate blastn searches against the two sets of reads. We selected the best hit for each CDS and noted the percent identity, alignment length, number of mismatches, number of gaps opened, the e-value, and bit score for each hit and calculated the mean values for both FY16936 and CBS5679.

To compare sequence identity between CBS5679 and FY16936 assemblies and a previously published *S. japonicus* var. *versatilis* DNA sequence (Naehring, Kiefer, & Wolf, 1995) we downloaded the Genbank sequence Z32848.1, encoding the 6.5 kb *S. japonicus* var. *versatilis* genes for 17S, 5.8S, and 25S ribosomal RNA (rRNA), which included the Internal Transcribed Spacer (ITS) 1 and ITS2. We generated BLAST databases for each genome and used Z32848.1 as the query sequence in a blastn search. We then identified the coordinates of the highest scoring hit from both CBS5679 and FY16936 assemblies and using the BEDTOOLS getfasta tools (v2.23.0) (Quinlan, 2014), we extracted those sequences from their respective genomes and used Muscle (v3.8.1551) (Edgar, 2004) to generate pairwise alignments between them and Z32848.1.

### Phylogenetic analyses

We downloaded publicly available (as of February 2023) genome assemblies of *Schizosaccharomyces* species as follows:

*S. octosporus* - GCA_000150505.2 (Tong et al., 2019)

*S. osmophilus* - GCA_027921745.1 (Jia et al., 2023)

*S. cryophilus* - GCA_000004155.2 (Tong et al., 2019)

*S. pombe* - PomBase (Harris et al., 2022)

*S. japonicus* (FY16936) - Ensembl Fungi GCA_000149845.2 (Yates et al., 2022)

*S. japonicus* (FY16936) – EI 1.0 GCA_956476325

*Saitoella complicata* (outgroup) - GCF_001661265.1 (Nishida, Hamamoto, & Sugiyama, 2011)

For each assembly we ran BUSCO (v5.4.4) using the most up-to-date Ascomycota database (ascomycota_odb10) to identify single-copy orthologs in each assembly. We then took the intersect of orthologs that were found to be complete and in a single copy across all assemblies and created a BED-format file of all those genes across each assembly. Using BEDTools getfasta tools (v2.23.0) (Quinlan, 2014) we extracted the sequence for each ortholog in each species. Next, we created ortholog-specific fasta files for each species and used Muscle to generate multispecies alignments for each ortholog.

We used IQTree2 (v2.2.2.2) (Minh et al., 2020) with default parameters (ModelFinder, tree search, ultrafast bootstrap (1000 replicates) and SH-aLRT test (1000 replicates)) to create a maximum likelihood (ML) phylogeny using all ortholog-specific gene alignments. Then, using Newick utils (v1.5) (Junier & Zdobnov, 2010) we set *S. complicata* as the outgroup. In order to obtain node support for the consensus phylogeny, we used IQTree2 to create individual ML phylogenies for each ortholog and then counted the number of times each branch in the original ML tree occurred in the set of ortholog-specific trees, using this rate as the bootstrapping values.

### Dating the CBS5679 split from the ‘type’ strain of S. japonicus

We used seqkit (W. Shen, Le, Li, & Hu, 2016) to concatenate the Muscle alignments for each ortholog. We then used SNP-sites (Page et al., 2016) to extract all variable sites where each nucleotide was either A, T, C, or G. Next, we used RelTree ML and RelTree Branch Lengths in MEGA (Stecher, Tamura, & Kumar, 2020; Tamura et al., 2012; Tamura, Tao, & Kumar, 2018) to date the CBS5679-FY16936 split, using the ortholog SNP-sites alignment data and the ML phylogeny. We used the following previously calculated divergence times (Jia et al., 2023; X. X. Shen et al., 2020):

Split 1. *S. japonicus* – *S. pombe* split (207.2 million years ago (MYA))

Split 2. *S. pombe* – *S. octosporus* split (108.2 MYA)

Split 3. *S. octosporus* – *S. cryophilus* split (29.4 MYA)

Split 4. *S. octosporus* – *S. osmophilus* split (15.7 MYA)

To compare our calculated divergence time with other previously calculated times, we ran RelTree ML and RelTree Branch Lengths, omitting Split 4 above (*S. octosporus – S. osmophilus*) and compared our predicted divergence time to that of the previously calculated divergence time (Jia et al., 2023).

### Genetic crosses

Crosses were performed on Sporulation Agar (SPA) medium (Munz & Leupold, 1970) with the following modifications: 3% agar, and 45 mg/ml each of adenine, histidine, leucine, and uracil. Homothallic and heterothallic strains (see Table 1) were mixed at a 1:10 ratio and incubated for 48 hours at 25°C. Cell mixtures were incubated for 1 hour at 37°C with 0.5% v/v glusulase (Perkin Elmer, NEE154001EA) to digest cell walls of vegetative cells and asci, followed by three washes with 20% ethanol, to ensure that no viable vegetative cells remain. For each cross, 63 random spores were dissected on Yeast Extract with Supplements (YES) agar plates (Petersen & Russell, 2016) using a Singer tetrad dissector (Singer Instruments).

**Table 1.** Strain list, genotypes and collection notes for genetic crosses between different variants of *S. japonicus*.

| Oliferenko Collection No. | Genotype | Notes |
| --- | --- | --- |
| SOJ1 | *S. japonicus* homothallic | Deposited in ATCC as ATCC10660; in H. Niki collection as NIG2008 |
| SOJ5709 | *S. japonicus* (?) homothallic | Deposited in Westerdijk Institute collection as CBS5679 |
| SOJ3657 | *S. japonicus gpd1-mNeonGreen:kanMX6* h- | Oliferenko collection; generated in the background of *S. japonicus* h- matsj-P2028 (NIG2028 in H. Niki collection) |

Plates were incubated at 30°C for 48 hours, replica plated to YES agar plates containing 75 μg/ml G418 (Sigma-Aldrich, G8168), and incubated at 30°C for a further 24 hours. Plates were scanned using ChemiDoc imager (Bio-Rad) before and after replica plating. The presence or the absence of G418 resistance was confirmed by re-streaking colonies on new G418-containing YES agar plates. Experiments were repeated three times. Statistical analyses were performed in Prism 9 (GraphPad).

### Identifying structural variants

To identify syntenic chromosomal regions between the CBS5679 and FY16936 assemblies, we used the nucmer tool in the MUMmer toolsuite (Delcher, Salzberg, & Phillippy, 2003) to align contigs of the CBS5679 assembly to the three chromosomes of FY16936, visualising the output in Dot (Nattestad, 2020). In order to further identify synteny, translocations, and rearrangements involving coding genes, we used Support Protocol 2 (‘Visualizing syntenies between genomes using BUSCO markers’) as set out by (Manni et al., 2021). In order to confirm any large genome rearrangements, we aligned CBS5679 ONT reads >50 kb back to the CBS5679 assembly and visualised those regions using IGV (Thorvaldsdottir, Robinson, & Mesirov, 2013) to check long-read coverage across each region. Additionally, we calculated mean coverage across each rearrangement (including 10kb up and downstream) and compared that to the mean coverage for the whole contig.

### GO term enrichment

We inputted the list of 446 genes located in the major translocated region into g:Profiler (Raudvere et al., 2019) and searched all known genes using the reference organism as ‘*S. japonicus*’, using the default significance threshold of p>0.05. g:Profiler is a web-based collection of tools that performs functional enrichment analyses of Ensembl gene lists. It maps the gene lists to known functional information sources and detects statistically significantly enriched biological processes and pathways. It is one of the few GO term enrichment tool that allows direct analysis of Ensembl Fungi genes.

### Assembly of a scaffolded chromosome-scale assembly

Using information gleaned from the previous analyses, we assembled a chromosome-scale assembly by using alignment information and ortholog placements, and orienting and joining contigs from CBS5679 relative to the FY16936 assembly and gap-filling with N’s. For the mitochondrial genome, we completed a self-vs-self blast search and trimmed redundancy from overlapping ends.

## Results

### Genome assembly and quality control

After assembling with Canu and carrying out three rounds of Pilon polishing, our assembly consisted of 46 contigs, totalling 16,207,024 bps. We compared the number of single-copy orthologs resolved in the assembly to those in the Ensembl Fungi *S. japonicus* assembly (GCA_000149845.2) and found that the number of single-copy, fragmented, duplicated, and missing orthologs only varied by a maximum of 0.3% (Figure 1). Additionally, we compared repeat content of CBS5679 to that of a telomere-to-telomere genome assembly of *S. japonicus* (FY16936; referred to as *S. japonicus* EI 1.0 assembly; GCA_956476325). We found that CBS5679 had 4% more repeat content than the *S. japonicus* EI 1.0 assembly and, like that assembly, most identified repeats belonged to the LTRs and Helitron-like transposons (Table 2). Genome annotation files for coding sequences, rRNA genes, tRNA genes, and repeats can be found in the Supplementary Data (Supplementary data S1).

**Table 2.** Comparison of assembly repeat content (%) by class between CBS5679 and FY16936 (EI 1.0). Abbreviations: SINE - Short interspersed nuclear elements; LTR - long terminal repeat; RC – rolling circle transposons (Helitron-like transposons).

| Repeat class | CBS5679 | EI 1.0 |
| --- | --- | --- |
| Nucleotides masked | 3,033,726 | 2,472,691 |
| % SINE | 0.004 | 0.10 |
| % LTR | 12.58 | 11.56 |
| % RC (Helitron) | 1.02 | 2.02 |
| % Unknown | 4.32 | 1.21 |
| % masked | 18.7 | 14.89 |

**Figure 1.**
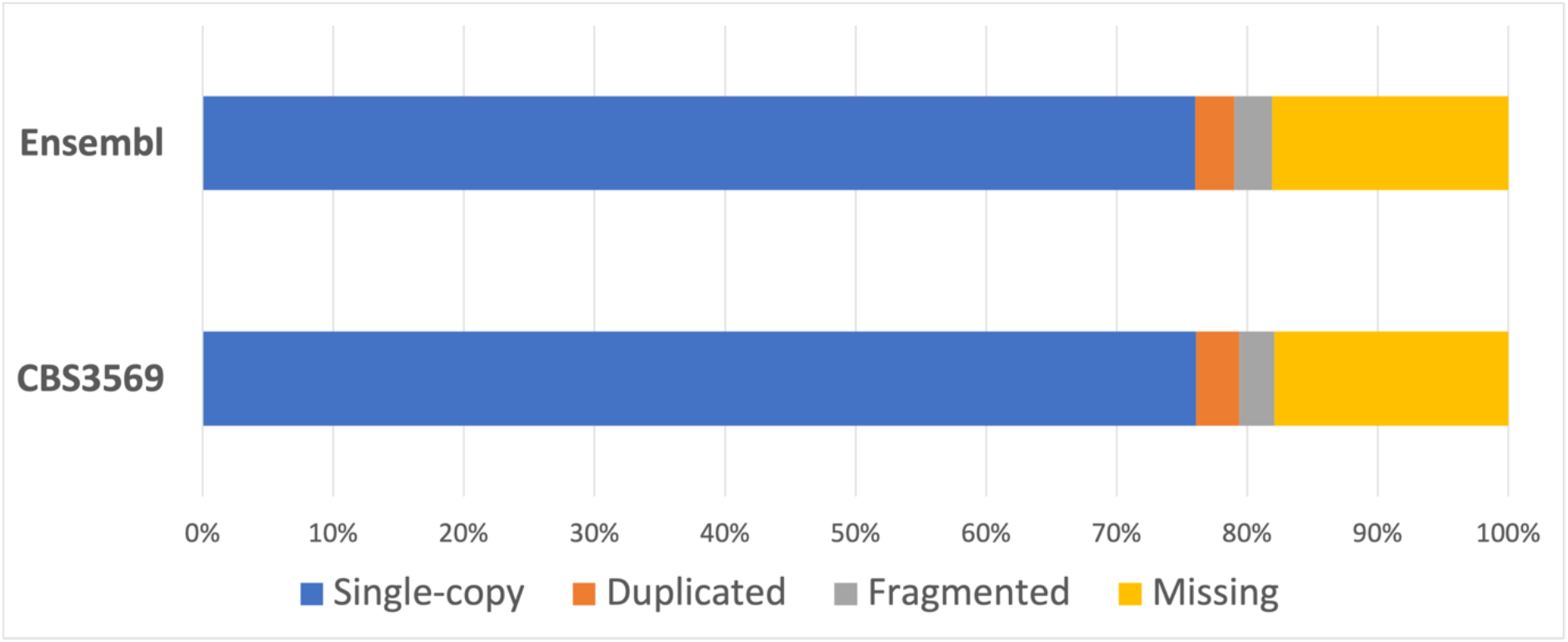
Comparison of BUSCO single-copy ortholog resolution between CBS3569 and Ensembl Fungi S. *japonicus*.

We compared the percentage nucleotide identity between Nanopore reads of FY16936 and CBS5679 to that of the CDS sequences of *S. japonicus* in Ensembl Fungi. We found that FY16936 reads showed a 99.96% identity to *S. japonicus* CDS sequences and CBS5679 showed only 88.44% identity (Table 3). This difference in identity could not be explained by Nanopore sequencing error rates (which would be equivalent for both datasets) or differences in mean alignment lengths (which are also similar). The number of mismatches and gaps opened in CBS5679 were 180 and 62 times greater (respectively) than in FY16936.

**Table 3.** Comparison of ONT read alignments for *S. japonicus* FY16936 and CBS5679 against CDS sequences for the Ensembl Fungi *S. japonicus* assembly.

| Dataset | % identity | Mean alignment length | Mean No. mismatches | Mean No. gaps opened |
| --- | --- | --- | --- | --- |
| FY16936 reads | 99.96 | 1693 | 0.098 | 0.25 |
| CBS5679 reads | 88.44 | 1641 | 176.036 | 15.39 |

### Comparison to S. japonicus var. versatilis rRNA

We compared 6.5 kb of previously reported sequence from the SSU, 5.8S, and LSU rRNA genes (including ITS1 and ITS2) from *S. japonicus* var. *versatilis* (Z32848.1) to that of CBS5679 and FY16936 (Table 4). We did not identify any differences within ITS1 or 5.8S. Across all features, *S. japonicus* var. *versatilis* and CBS5679 were the most similar (25 SNPs), followed by CBS5679 and *S. japonicus* EI 1.0 (48 SNPs). Finally, *S. japonicus* var. *versatilis* and *S. japonicus* EI 1.0 showed the greatest distance (73 SNPs).

**Table 4.** Pairwise comparisons of 17S, 5.8S, and 25S ribosomal RNA genes, including Internal Transcribed Spacer (ITS) 1 and ITS2 between *S. japonicus* var. *versatilis*, CBS5679, and *S. japonicus* EI 1.0. Values represent divergence, calculated by the number of SNPs divided by the length of the feature. Numbers in brackets are the absolute number of SNPS.

| Feature | length | *S. japonicus* var. *versatilis* vs CBS5679 | *S. japonicus* var. *versatilis* vs EI 1.0 | CBS5679 vs EI 1.0 |
| --- | --- | --- | --- | --- |
| 17S | 2019 | 0.00149 (3) | 0.00198 (4) | 0.00050 (1) |
| ITS1 | 186 | 0 | 0 | 0 |
| 5.8S | 157 | 0 | 0 | 0 |
| ITS2 | 268 | 0 | 0.08582 (23) | 0.08582 (23) |
| 25S | 3433 | 0.00641 (22) | 0.01340 (46) | 0.00699 (24) |
| Total | 6063 | 0.00412 (25) | 0.01204 (73) | 0.00792 (48) |

### *Phylogenetic analysis of* CBS5679

We ran BUSCO on CBS5679 along with six other *Schizosaccharomyces* assemblies plus one Taphrinomycotina outgroup (*Saitoella complicata*) and resolved 1023 single-copy orthologs present in all assemblies. After creating ortholog-specific alignments, we generated a ML tree using all alignments. The best-fit model of GTR+F+I+R2 was chosen according to the Bayesian information criterion (BIC). We then created ML trees for each of the 1023 orthologs and used those trees to evaluate support for the original ML tree (Figure 2). CBS5679 splits from both *S. japonicus* assemblies with 99% bootstrap support, providing high confidence for the split.

**Figure 2.**
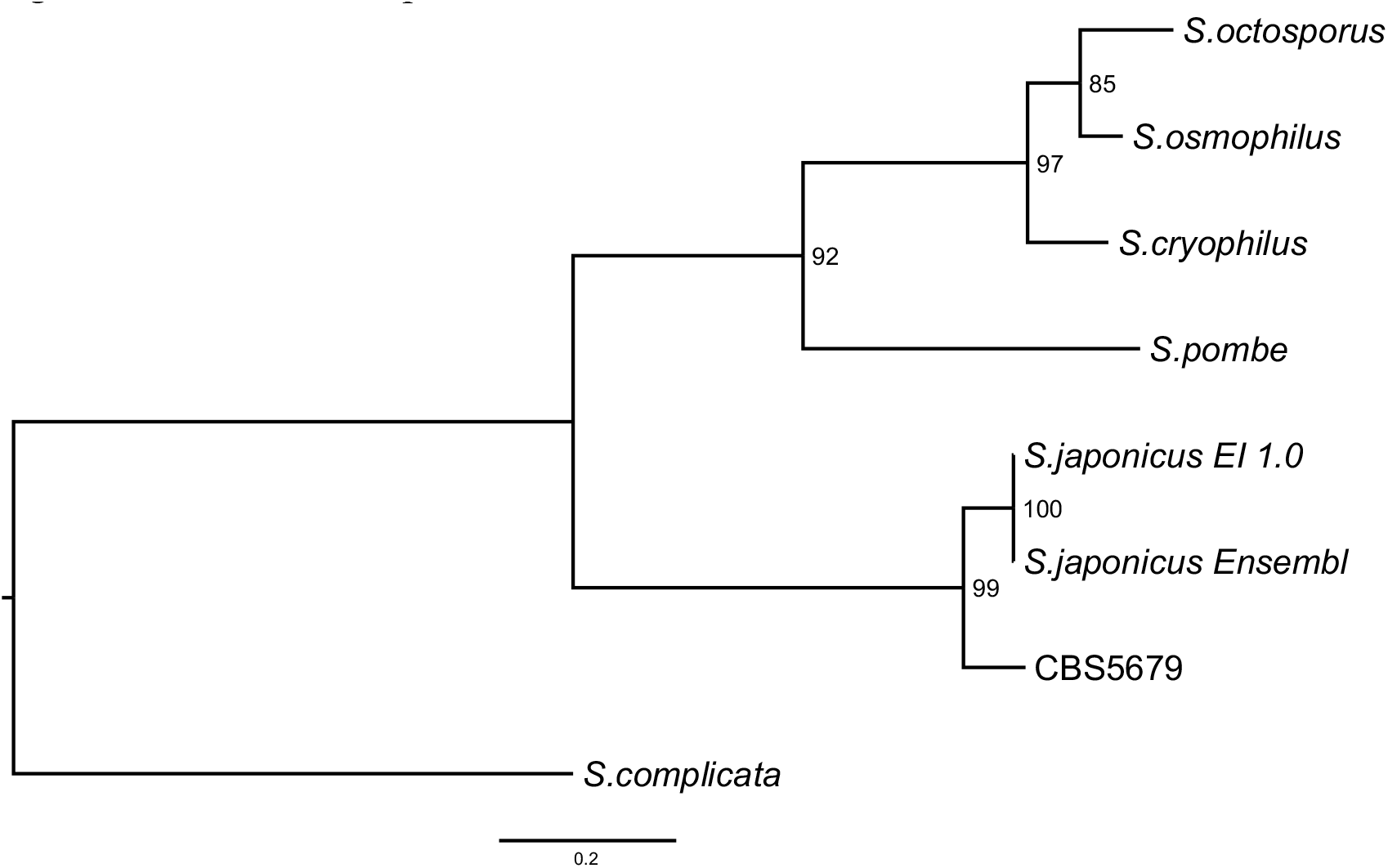
Maximum likelihood (ML) tree based on 1023 single-copy orthologs. Node labels represent support values from individual ML ortholog trees.

### CBS5679 has split from S. japonicus approximately 25 million years ago

Using the ML tree generated above along with 968,591 variable sites contained within the 1023 concatenated, aligned single-copy orthologs, and previously calculated divergence times, we calculated the divergence time of *S. japonicus* and CBS5679. Using RelTree ML, the split was dated at 25.05 MYA (Figure 3) and using RelTree branch length, 24.69 MYA. Additionally, we removed the previously calculated split between *S. octosporus* and *S. osmophilus* from the tree to confirm that this dating-method recapitulated the previously calculated divergence time (Jia et al., 2023). RelTree ML and RelTree branch lengths dated this split at 16.27 MYA and 16.02 MYA respectively (and confirmed the *S. japonicus* and CBS5679 divergence time), closely reflecting the previously calculated divergence time of 15.7 MYA (Jia et al., 2023) (Supplementary Figure S1).

**Figure 3.**
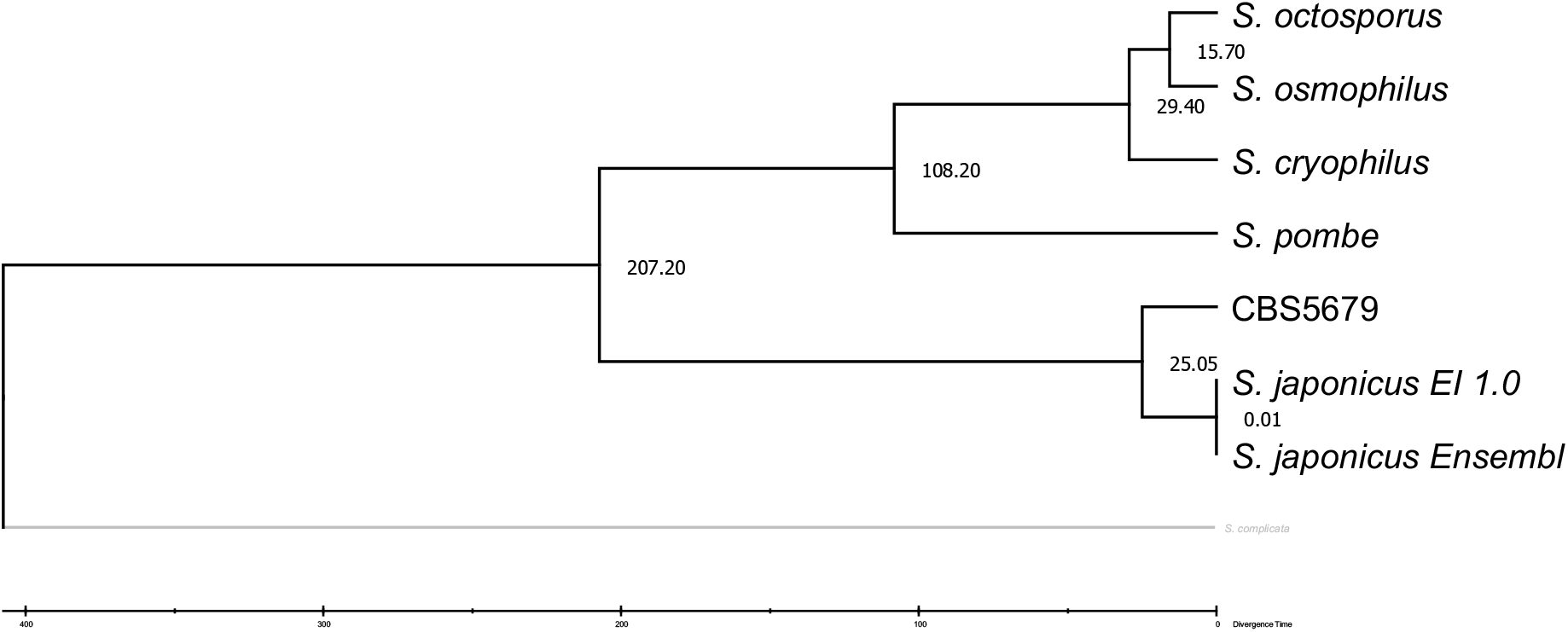
Dated divergence of CBS5679 from *S. japonicus* using RelTree ML. The node numbers represent divergence times in millions of years.

### CBS5679 does not generate viable progeny in crosses with the type S. japonicus strain

To test if the CBS5679 strain (Bouwknegt et al., 2021) could generate viable progeny with *S. japonicus*, we performed crosses between the homothallic CBS5679, or the “type” homothallic *S. japonicus* isolate (ATCC10660) and a heterothallic h-strain of *S. japonicus* (Furuya & Niki, 2009), where the glycerol-3-phosphate dehydrogenase Gpd1 was marked with the G418-resistance cassette (Ying Gu, Alam, & Oliferenko, 2023) .We used the heterothallic h-strain in excess (see Materials and Methods for details), since both homothallic strains sporulate with high efficiency. As controls, we included either CBS5679 or ATCC10660 sporulating individually.

Following sporulation, we dissected and plated individual spores to ensure that no vegetative cells contribute to our analyses. Spores originating from crosses containing the CBS5679 strain germinated and formed colonies at higher frequency as compared to those containing the “type” ATCC10660 *S. japonicus* isolate (Figure 4A). The introduction of the heterothallic h-strain into crosses with either homothallic strain did not affect overall spore germination and colony formation efficiency (Figure 4A). Of note, we observed the appearance of G418-resistance only in the crosses between ATCC10660 and the G418-resistance-marked h-*S. japonicus* strain (Figure 4B). In our experimental conditions, the proportion of G418-resistant colonies was consistent between three different experiments and averaged ∼34%. We concluded that it was likely that CBS5679 strain has become reproductively isolated from the “type” strain of *S. japonicus*.

**Figure 4.**
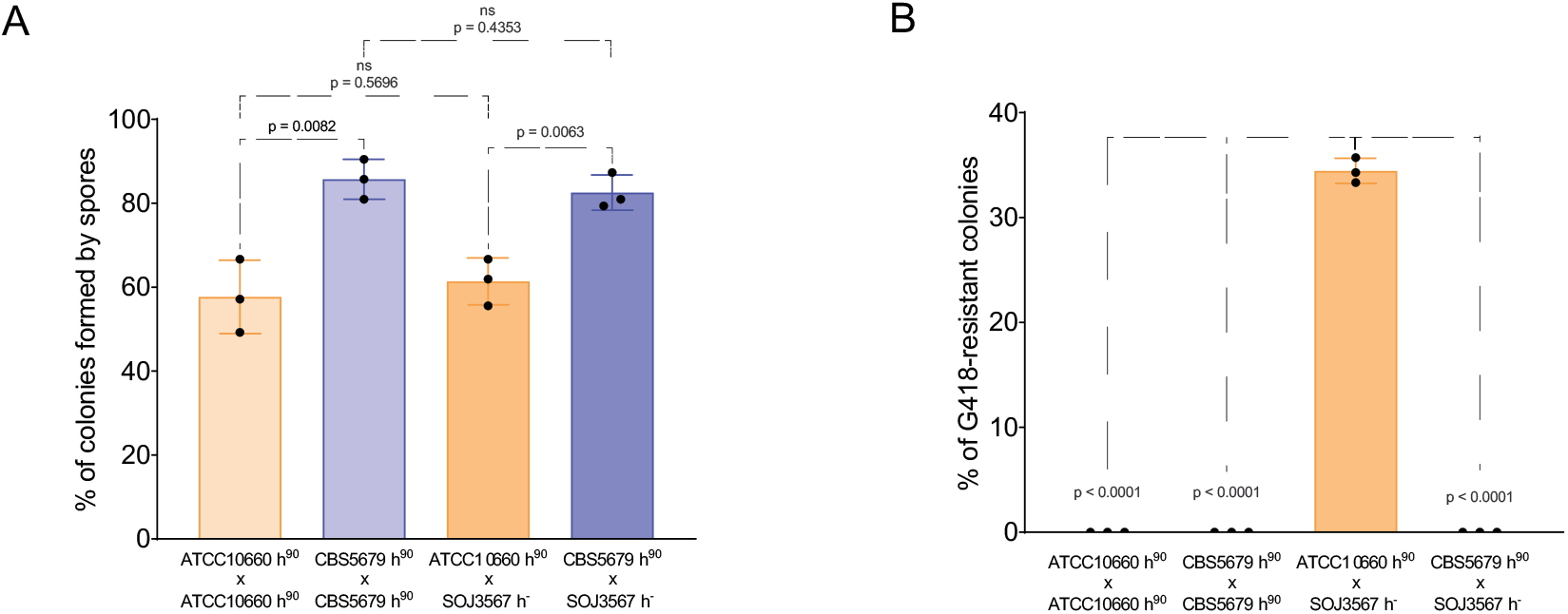
(A) Graph illustrating the efficiency of colony formation by spores formed in crosses of indicated genotypes. (B) Graph showing the percentage of G418-resistant colonies in crosses of indicated genotypes. (A-B) Experiments were performed in three biological replicates and *p-values* were derived using unpaired parametric t test.

### The genomes of CBS5679 and the type S. japonicus strain exhibit major structural variation

Using Nucmer, we aligned the contigs from CBS5679 to that of the three chromosomes of *S. japonicus* EI 1.0 to identify syntenic regions, translocations, and inversions (Figure 5). Additionally, we plotted the BUSCO single-copy orthologs present on both genomes to identify the genomic locations of orthologs between the two assemblies (Figure 6).

**Figure 5.**
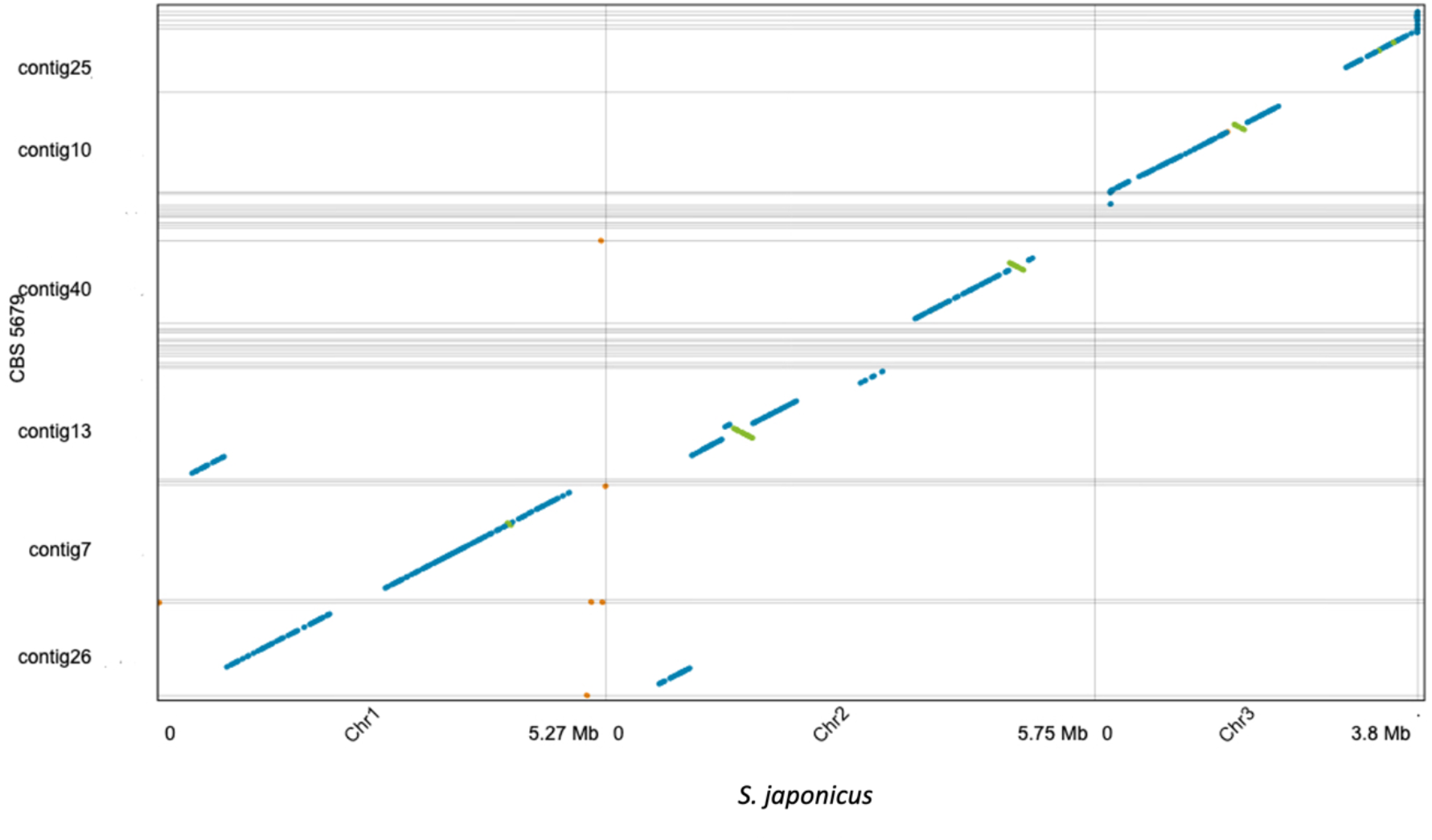
Alignment of CBS5679 to *S. japonicus* EI 1.0 showing syntenic regions and translocation involving Chr1 and Chr2. Blue lines refer to forward alignments and green lines refer to reverse alignments (inversions). Reciprocal translocations can be seen between *S. japonicus* ‘Chr1’ and ‘Chr2’ and CBS5679 ‘contig13’ and ‘contig26’.

**Figure 6.**
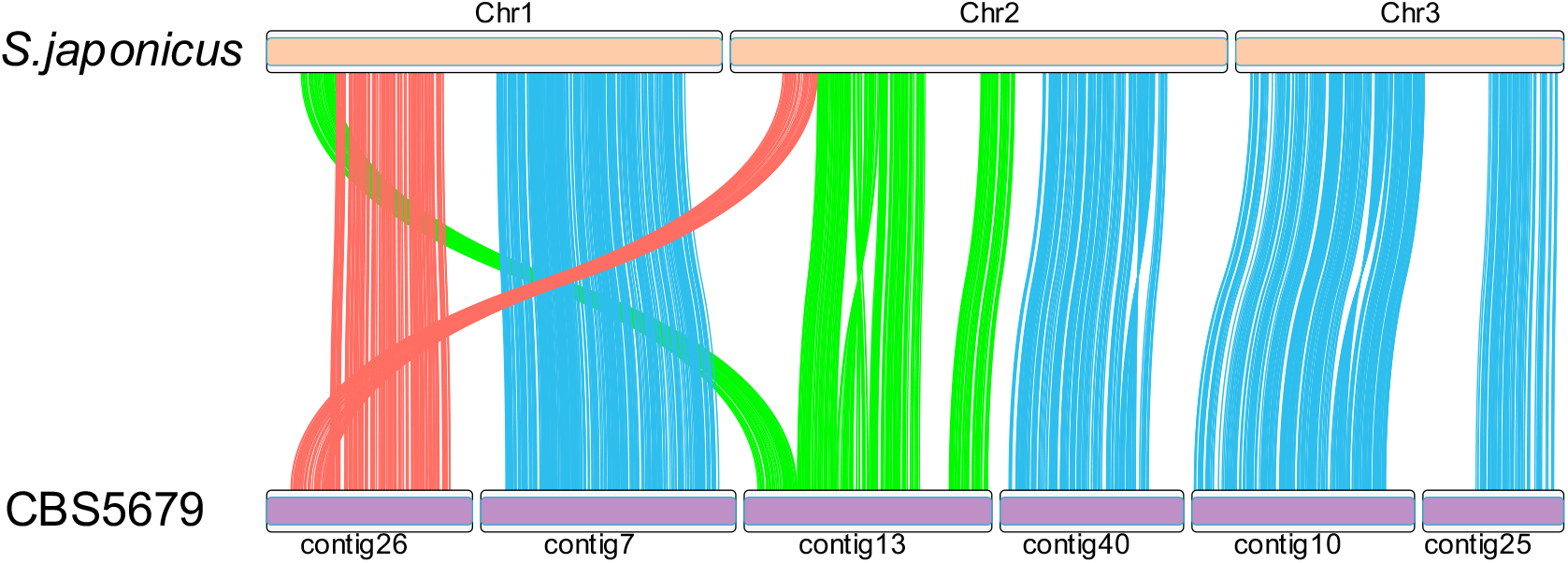
Placement of BUSCO single-copy orthologs present on both *S. japonicus* EI 1.0 and CBS5679. Orthologs present on CBS5679 contig13 are mapped in green and those on contig26 are mapped in red, showing the location of orthologs involved in a reciprocal translocation.

It is clear there is a reciprocal translocation involving chromosomes 1 and 2 in *S. japonicus*, and contig26 and contig13 on CBS5679. Using the mapping locations of the orthologs (Supplementary Table S1), we identified the approximate location of the translocations, which occurred between position 799,696 and 808,427 on Chr 1 and positions 1,004,860 and 1,010,039 on Chr 2 (Table 5).

**Table 5.** The approximate sites of the translocation between *S. japonicus* Chr 1 and Chr2 and CBS5679 contig13 and contig26. The ortholog name is that from BUSCO odb10 and the co-ordinates refer to each given ortholog in each assembly.

|  | *S. japonicus* EI 1.0 |  |  | CBS5679 |  |  |
| --- | --- | --- | --- | --- | --- | --- |
| Ortholog name | Chromosome | Ortholog start | Ortholog end | Contig | Ortholog start | Ortholog end |
| 85315at4890 | 1 | 797053 | 799696 | contig13 | 536839 | 539492 |
| 213148at4890 | 1 | 808427 | 810400 | contig26 | 1457191 | 1459164 |
| 338471at4890 | 2 | 1004387 | 1004860 | contig26 | 1464423 | 1464842 |
| 202577at4890 | 2 | 1010039 | 1011033 | contig13 | 547711 | 548705 |

We aligned CBS5679 ONT reads >50kb back to the CBS5679 assembly and visualised the regions on contig13 and contig26, which contained the translocations (Supplementary Figure S2). In both cases it was clear that no misassemblies had taken place with long reads spanning both regions. Additionally, mean long-read coverage across each translocated region of CBS5679 was close to the mean for the whole contig. Specifically, the mean coverage for contig26 was 10 (SD 3.25) and the mean coverage over the translocation was 10 (SD 0.66). The mean coverage for contig13 was 9 (SD 4.23) and the mean coverage over the translocation was 7 (SD 1.42).

In addition, three large inversions can be seen between the two assemblies (denoted by the green lines in Figure 5, and large ‘twists’ in the ortholog mapping in Figure 6). The length of the large inversions varied between 147.2 kb and 307 kb (Table 6).

**Table 6.** Location and size of inversions between CBS5679 and *S. japonicus* EI 1.0. Locations are given relative to EI 1.0. CBS5679 contig refers to the contig containing the inversion.

| EI 1.0 chromosome | start | end | size | CBS5679 contig |
| --- | --- | --- | --- | --- |
| 2 | 1416577 | 1723660 | 307,084 | contig13 |
| 2 | 4763199 | 4960096 | 196,898 | contig40 |
| 3 | 1615898 | 1763107 | 147,210 | contig10 |

### Major chromosomal rearrangement carries genes associated with metabolism and cellular organization

Large genomic rearrangements are often responsible for pre-zygotic barriers between interspecies hybridisation. We identified genes on translocated genomic loci and carried out Gene Ontology (GO) term enrichment analyses. Given a list of genes from an experiment, GO term enrichment identifies the pathways each gene is involved in and then identifies pathways that have more genes than expected by chance. We identified 446 protein-coding genes situated in chromosomes 1 and 2 that lay within the reciprocal translocation with CBS5679 and analysed them for enriched GO terms (Table 7). Although many of the significant terms were high-level (e.g., ‘molecular_function’, ‘biological_process’, ‘cellular_process’, etc.), many other terms fell into one of two different terms: “metabolic processing”, and “cellular assembly”. Metabolic processing terms include homoserine metabolic process, primary metabolic process, organic substance metabolic process, and metabolic process, whilst cellular assembly terms include cellular component assembly, cellular component biogenesis, cellular anatomical entity, cellular component, actin cortical patch, endocytic patch, and cortical actin cytoskeleton.

**Table 7.** Enriched gene ontology (GO) terms identified from 446 protein-coding genes located within the reciprocal translocation. The GO categories are as follows: GO:MF – molecular function, GO:BP – biological process, GO:CC – cellular component. The full analyses, including all input data, run parameters and options may be found at the following URL: https://biit.cs.ut.ee/gplink/l/DclCMvGJSh.

| GO category | Term name | GO term ID | Adjusted p-value |
| --- | --- | --- | --- |
| GO:MF | molecular_function | GO:0003674 | 7.61E-08 |
| GO:BP | biological_process | GO:0008150 | 1.48125E-05 |
| GO:BP | cellular process | GO:0009987 | 0.000756756 |
| GO:BP | cellular component assembly | GO:0022607 | 0.001871718 |
| GO:BP | primary metabolic process | GO:0044238 | 0.002338023 |
| GO:BP | homoserine metabolic process | GO:0009092 | 0.012502301 |
| GO:BP | cellular component biogenesis | GO:0044085 | 0.017597583 |
| GO:BP | organic substance metabolic process | GO:0071704 | 0.024054927 |
| GO:BP | metabolic process | GO:0008152 | 0.030869142 |
| GO:CC | cellular anatomical entity | GO:0110165 | 2.56903E-06 |
| GO:CC | cellular_component | GO:0005575 | 9.65011E-06 |
| GO:CC | actin cortical patch | GO:0030479 | 0.017274955 |
| GO:CC | endocytic patch | GO:0061645 | 0.022400654 |
| GO:CC | cortical actin cytoskeleton | GO:0030864 | 0.046312161 |

### Assembly of a scaffolded chromosome-scale assembly

Our final assembly, (CATQFW010000000), comprised three autosomal chromosomes, one mitochondrial genome, plus 39 unplaced contigs. The combined total of the three autosomal chromosomes plus mitochondrial genome was 13.3Mb (including Ns) and 16.5 Mb including the unplaced contigs.

## Discussion

We have assembled and annotated the genome of CBS5679 that was originally assumed to be a strain of *Schizosaccharomyces japonicus* var. *versatilis*. The unscaffolded assembly is 16.2 Mb of which 12.9 Mb is represented by 6 contigs that each align to a chromosome arm of the reference strain of *S. japonicus*. The number of single-copy orthologs is almost identical to that of the Ensembl Fungi reference of *S. japonicus* and the percentage of the genome masked is similar, although the number of repeats of ‘Unknown’ class was greater in CBS5679. Yet, at the CDS level CBS5679 significantly diverges from S. *japonicus*, sharing only 88.4% sequenced identity. Phylogenetically CBS5679 is a separate sister clade to *S. japonicus* (with 99% support values). The timing of this split is estimated to be around 25.05 MYA, and the accuracy of this estimate is further supported by our estimate of the *S. octosporus* - *S. osmophilus* split, which was within 0.3 MY of the previously calculated split (Jia et al., 2023).

Over the LSU, 5.8S and SSU rRNA unit, CBS5679 shows much greater similarity to *S. japonicus* var. *versatilis*, showing approximately half as many differences to that strain than FY16936 (var. *japonicus*) and possibly reflects the genetic variation found within *S. versatilis*, or perhaps represents population structure within the species.

When compared to *S. japonicus* FY16936, CBS5679 has a reciprocal translocation between chromosomes 1 and 2, with roughly 800 kb from chromosome 1 on FY16936 now being located on CBS5679 chromosome 2, and 1 Mb of chromosome 2 on FY16936 now being located on CBS5679 chromosome 1. Additionally, CBS5679 contains at least three large scale (> 140 kb) inversions. In *Saccharomyces cerevisiae* it was found that if two genes were closely located, they were more likely to be involved in the same biological process (Santoni, Castiglione, & Paci, 2013). The largest metabolic gene cluster in yeast (the DAL cluster), was assembled through an almost simultaneous set of telomeric rearrangements that brought together the genes in this pathway (Wong & Wolfe, 2005). Furthermore, work on *S. pombe* showed that chromosomal rearrangements contribute to reproductive success and could be strongly beneficial in one environment but deleterious in another. These rearrangements were also accompanied by alterations in gene expression (Teresa Avelar, Perfeito, Gordo, & Godinho Ferreira, 2013). These large chromosomal rearrangements reported here could be one of the reasons for reproductive isolation between CBS5679 and the ‘type’ strain of *S. japonicus*.

Experimentally, we demonstrate that CBS5679 is reproductively isolated from the *S. japonicus* ‘type’ strain. A heterothallic h-strain of *S. japonicus*, marked with an G418-resistance gene, efficiently generated G418 resistant spores when crossed with a homothallic *S. japonicus* isolate, but not with the homothallic CBS5679 isolate. This cannot be explained by a failure in sporulation or germination for the CBS5679 isolate, since both processes were equivalent or occurred more efficiently in CBS5679 than in the *S. japonicus* ‘type’ strain.

Interestingly, GO enriched terms for genes located in the translocated regions fell mainly into two categories: metabolic processing, and cellular assembly. CBS5679 and other *S. versatilis* strains share a number of metabolic characteristics with the type strain of *S. japonicus*, including anaerobic growth and the inability to utilize non-fermentable carbon sources (Alam, Gu, Reichert, Bahler, & Oliferenko, 2023; Bouwknegt et al., 2021; Bulder, 1963, 1971; Wickerham & Duprat, 1945). Yet, some aspects of amino acid biosynthesis and energy and redox metabolism might have been rewired after the split between the two species. For instance, it would be of interest to revisit the older biochemical data on the differences in the complement of cellular cytochromes between *S. japonicus* and *S. versatilis* in light of the newly available genomes and our current understanding of fission yeast metabolism and physiology (Alam et al., 2023; Ying Gu et al., 2023; Kaino, Tonoko, Mochizuki, Takashima, & Kawamukai, 2018).

Within the cellular assembly category, cortical actin and endocytic patch terms are significantly enriched. Cortical actin filaments nucleated by formins mediate long-range vesicle trafficking, which is required for the polarized growth of vegetative cells and the formation of specialized fusion structures essential for mating. A different type of actin networks, the Arp2/3-nucleated “patches” promote the formation and movement of endocytic vesicles, enabling nutrient uptake and polarized growth. During cell division, actin-based structures ensure cortical ingression and eventual separation of the daughter cells (Kovar, Sirotkin, & Lord, 2011; Mishra, Huang, & Balasubramanian, 2014). Together with metabolism, actin cytoskeleton impinges on virtually every aspect of biology. Understanding the phylogenetic relationship between *S. japonicus* and *S. versatilis* and the availability of high-quality genome assemblies for both species will undoubtedly facilitate mechanistic evolutionary cell biology studies using fission yeasts.

CBS5679 diverged from *S. japonicus* around 25 MYA and is only 88% identical at the CDS level. Its genome contains large-scale translocations and inversions, and it does not readily cross with *S. japonicus* var. *japonicus*. Thus, we conclude that CBS5679 represents a distinct species. CBS5679, originally listed as *S. japonicus* var *versatilis* (Phaff et al., 1964), shows the closest identity to CBS103, the type strain of *S. japonicus* var *versatilis* (Wickerham & Duprat, 1945). Further work comparing CBS103 and CBS5679 is required to identify if the differences observed between them indeed represents intra-species diversity, or if CBS5679 constitutes a closely related separate species. The CBS103 and CBS5679 strains have been deposited in databases as *S. japonicus var versatilis*. Our work suggest that the taxonomic status of *S. japonicus* var. *versatilis* is raised and that it should be treated as a separate species within the fission yeast clade, *Schizosaccharomyces versatilis*.

## Acknowledgements

We acknowledge Frank Uhlmann for reading and commenting on this manuscript. We thank Li-Lin Du and Guo-Song Jia for helpful comments and sharing unpublished results.

GJE assembled and annotated the assembly and carried out all bioinformatic and phylogenetic analyses. EGG and SO carried out the genetic crosses. CN, SO, and WH assisted with experimental design. All authors contributed to writing and reviewing the manuscript.

## Data availability

The genome assembly for *S. versatilis* can be found under ENA Project PRJEB63472, with an accession number of CATQFW010000000.

## Supplementary Data

**Supplementary Figure S1.**
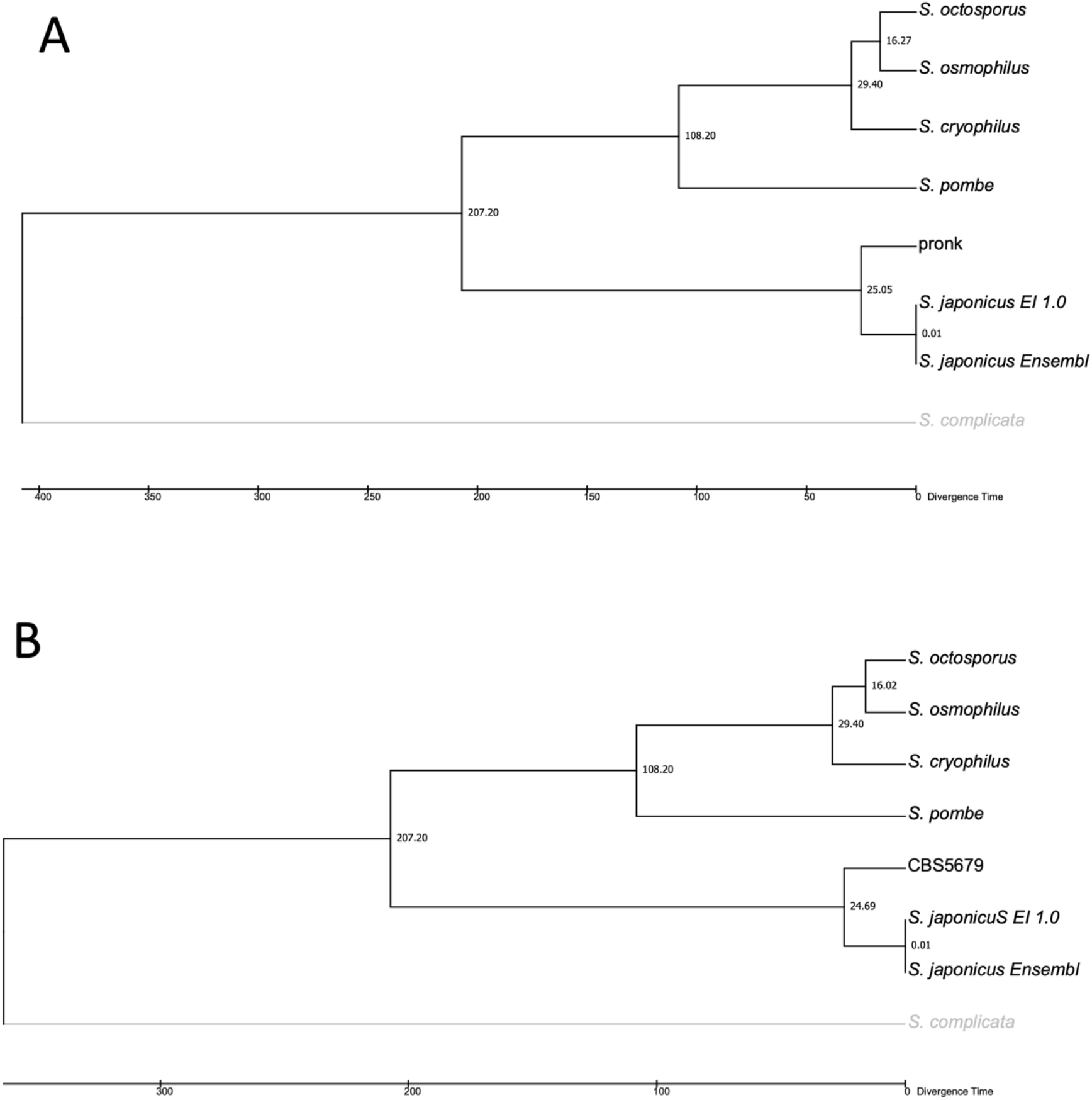
Calibration of CBS5679 - *S. japonicus* split using (A) RelTree Maximum Likelihood and (B) RelTree Branch Lengths. The node numbers represent divergence times in millions of years. In the trees, the previously-calculated timing of the *S. octosporus – S. osmophilus* split (15.7 MYA) was removed and recalculated along with the *S. japonicus –* CBS56679 split, confirming our previously calculated divergence time of 25 MYA.

**Supplementary Figure S2.**
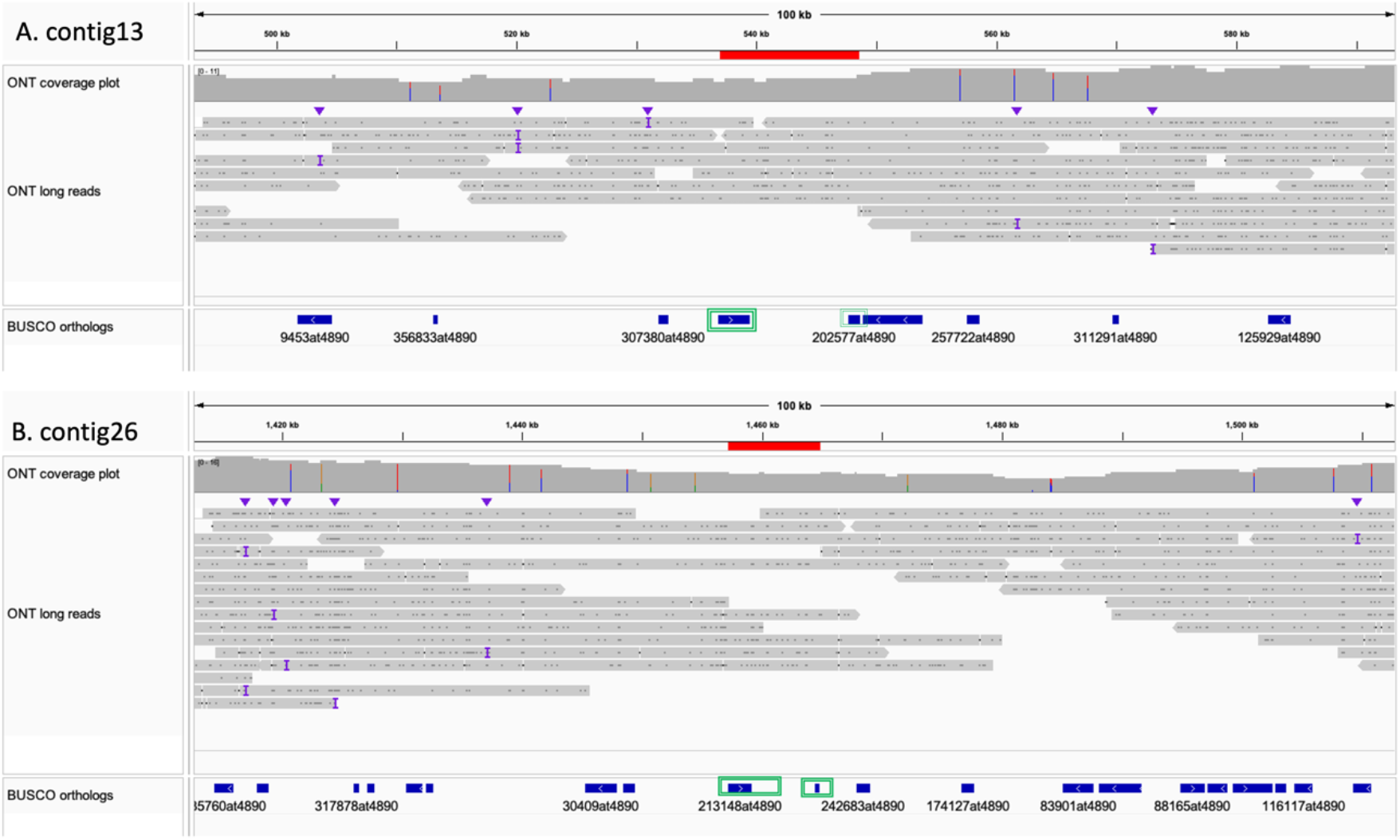
Long-read (ONT) coverage across translocations on (A) contig13 and (B) contig26, using the IGV genome browser. The solid red line represents the translocations (relative to *S. japonicus* chromosomes 1 and 2), the green squares highlight the flanking BUSCO orthologs, and the grey horizontal bars in the ‘ONT long reads’ panel represent individual ONT reads across the translocations.

Supplementary Table S1

Co-ordinates of the BUSCO single-copy orthologs present in both CBS5679 and *S. japonicus* EI 1.0. The table is sorted by the co-ordinates of orthologs for the chromosome-scale assembly *S. japonicus* EI 1.0.

Supplementary Data S1 (s_versatilis.gff.tar.gz)

This archive contains four annotation files in GFF format:

s_versatilis_liftoff.gff – Genome annotation from Ensembl Fungi assembly of *S. japonicus* using Liftoff (Shumate & Salzberg, 2020).

s_versatilis_repeats.gff – Annotation of repeat content from RepeatModeler v2.0.3 (Flynn et al., 2020) and RepeatMasker (v4.1.2-p1) (Smit et al., 2015; Tarailo-Graovac & Chen, 2009). s_versatilis_trnascan.gff – Annotation of tRNA loci using tRNAscan-SE (v2.0.12) (Chan et al., 2021)

s_versatilis_barrnap_rrna.gff – Annotation of rRNA loci using barrnap (v0.9) (Seeman, 2018).

